# Dissociable Neural Systems Support the Learning and Transfer of Hierarchical Control Structure

**DOI:** 10.1101/2020.01.16.909978

**Authors:** Adam Eichenbaum, Jason M. Scimeca, Mark D’Esposito

## Abstract

Humans can draw insight from previous experiences in order to quickly adapt to novel environments that share a common underlying structure. Here we combine functional imaging and computational modeling to identify the neural systems that support the discovery and transfer of hierarchical task structure. Human subjects completed multiple blocks of a reinforcement learning task that contained a global hierarchical structure governing stimulus-response action mapping. First, behavioral and computational evidence showed that humans successfully discover and transfer the hierarchical rule structure embedded within the task. Next, analysis of fMRI BOLD data revealed activity across a frontal-parietal network that was specifically associated with the discovery of this embedded structure. Finally, activity throughout a cingulo-opercular network and in caudal frontal cortex supported the transfer and implementation of this discovered structure. Together, these results reveal a division of labor in which dissociable neural systems support the learning and transfer of abstract control structures.

## Introduction

Whether it is learning how to drive a new car, interacting with an unfamiliar social group, or navigating a foreign airport, humans show remarkable adaptability by inferring the correct action given minimal information. Such learning usually occurs via trial and error, where the individual updates their behavior based on feedback received after each action. Even though trial and error learning can be slow, humans routinely accelerate their problem-solving approaches by generalizing knowledge from the past. Indeed, the ability to generalize from past experiences has been an area of interest for cognitive scientists for well over a century (Woodworth & Thorndike, 1901). The process by which prior experiences can generalize to behavior in a novel context is generally referred to as “transfer” in the cognitive sciences, and “inductive bias” in machine learning (Botvinick et al., 2019; Tervo, Tenenbaum, & Gershman, 2016).

When simple stimulus-response mappings are learned across multiple task blocks, responses learned in one context can be directly transferred to a subsequent context, such that a performance benefit can be immediately observed (Behrens, Woolrich, Walton, & Rushworth, 2007; Collins, Cavanagh, & Frank, 2014; Collins & Frank, 2016). Although humans can encounter scenarios such as these (e.g., opening computer applications on a Windows vs. Apple operating system), humans can also find themselves in settings where directly transferring previously rewarded behaviors results in failure (e.g. starting computer programs on Windows/Apple vs. using the command line on Linux). In cases where direct transfer of previously learned responses is not feasible, it is advantageous to leverage a more abstract form of transfer where instead prior knowledge is used to guide the learning of the correct behavior in a novel context, a learning process known as “learning to learn” (Bavelier, Green, Pouget, & Schrater, 2012; Botvinick et al., 2019; Harlow, 1949; Kemp, Goodman, & Tenenbaum, 2010). While the behavioral and neurobiological underpinnings of more direct types of transfer have been relatively well-characterized (Collins et al., 2014; Collins & Frank, 2016), the neural systems and mechanisms underlying this more abstract form of transfer remain poorly understood.

Everyday experiences are often structured hierarchically, in that actions and experiences are influenced by a series of superordinate contexts and rules. For example, when traveling away from home it is common to pack a bag with clothes and overnight necessities. However, the rule that restricts packing small-volume liquids is only relevant in certain contexts: when traveling by airplane, but not by car. By grouping these sets of behaviors and experiences in a hierarchical fashion, one is able to easily generalize rules from one context to another, and even to contexts that have not yet been personally experienced. If the restriction of liquids is due to a rule associated with airport security, then one can extrapolate that modes of transportation that do not use airport security, even those that they have not yet taken (e.g., boat, train, etc.), will not be subject to liquid restrictions.

One way in which a learned hierarchical structure may be generalized to novel contexts is the creation of task sets or task structures that span across related contexts regardless of low-level features (Collins & Frank, 2013). For example, an individual could have a specific task set for their home airport, one that incorporates information about where the security line forms, which terminal each airline uses, and so on. However, it is advantageous to have a superordinate task set that spans airports in general, one that abstracts away from specific features of each airport and instead contains shared high-level details, such as the general spatial arrangement: ticketing booths are in front of the security checkpoint, while the shopping concourse and gates are behind security. Such a superordinate structure would allow for previous information to guide new action in a specific environment that the individual has never previously encountered (e.g, a foreign airport).

Although the combination of task sets and hierarchical processing provides a natural candidate solution for how learned hierarchical structure is generalized to new contexts, the neural basis of these cognitive processes has typically been studied in isolation. Growing neurobiological and computational evidence suggests that the frontal cortex facilitates the learning of complex, hierarchically structured behavior (Badre & D’Esposito, 2007; Badre, Kayser, & D’Esposito, 2010; Badre & Nee, 2018; Collins & Frank, 2013; Frank & Badre, 2012; Koechlin, 2003; Nee & D’Esposito, 2016; Wang et al., 2018). Regions along lateral frontal cortex that support such behavior are functionally organized along a rostrocaudal gradient, such that increasingly rostral regions resolve competition between available motor responses in accord with the level of task abstraction (Badre & D’Esposito, 2007). Processing of task sets has generally been related to activity in frontal cortex, as well as to a distributed network of regions referred to as the “cingulo-opercular network” (Dosenbach, Fair, Cohen, Schlaggar, & Petersen, 2008; Sakai, 2008).

In the current experiment, we designed a hierarchical reinforcement learning task that promotes the creation and transfer of a superordinate structure (Figure 1). Subjects learned stimulus-response associations in multiple blocks for which hierarchical task structure existed. Specifically, a 2^nd^-order hierarchical rule determined that the shape of the stimulus cued 1^st^-order rules defined by other stimulus dimensions (e.g. if the stimulus is a square, perform action 1 for red squares, and action 2 for blue squares, however if the stimulus is a circle, perform action 3 for striped circles, and action 4 for checkered circles, regardless of other stimulus features). Critically, although each block contained entirely new stimulus features, the 2^nd^-order hierarchical rules (hereafter referred to as 2^nd^-order policy) remained. Therefore, successful performance of a previous block conveyed no advantage to immediate performance on subsequent blocks. However, knowledge of the correct hierarchical policy structure would instead facilitate a more rapid learning of the correct mappings. Specifically, subjects learned stimulus response associations across five blocks wherein the first block contained no hierarchical policy structure, while the final four blocks all shared the same global hierarchical policy structure. Thus, subjects who learn the block-specific hierarchical policy in successive blocks can discover the existence of the global hierarchical structure. By transferring their knowledge of the global hierarchical structure to subsequent blocks, subjects can more rapidly learn the block-specific hierarchical policy. This design allowed us to show that subjects were capable of discovering and transferring the global hierarchical policy structure. In addition, we leveraged converging computational modeling approaches to examine the degree to which subjects transfer learned structure and how this transfer evolved over the course of the task. Lastly, we used fMRI to investigate the neural systems that support the initial discovery of global hierarchical policy structure and subsequent transfer of this structure knowledge to facilitate future learning.

**Figure 1.**
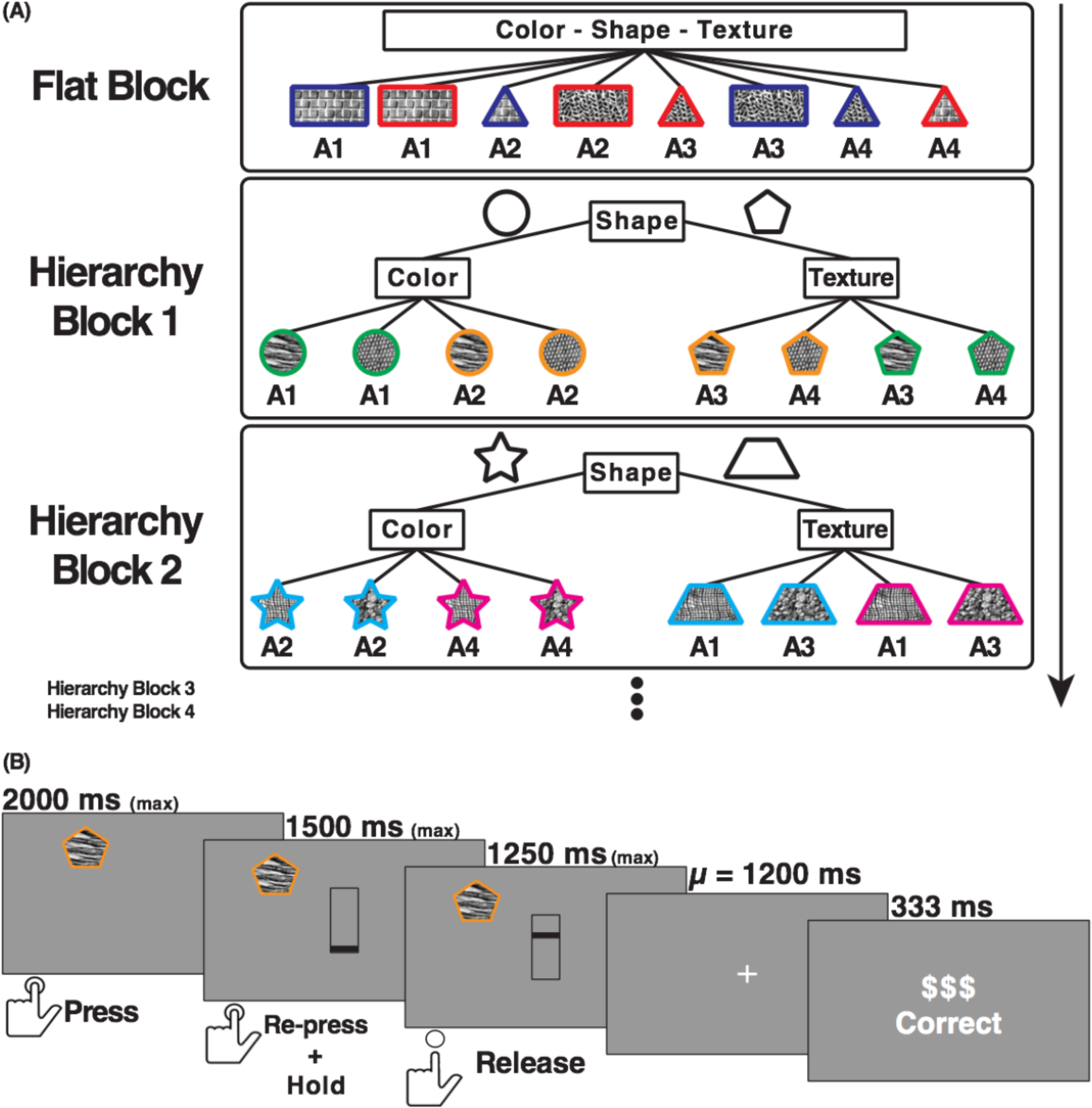
Schematic Depiction of Experimental Logic and Trial Sequence. (A) Schematic of task design showing example stimulus-to-action mappings. Subjects completed five blocks in total throughout the experiment. The stimuli in each block varied along three dimensions: shape, color, and texture. Each block contained two stimulus features for each dimension (e.g. two shapes) and the specific features changed for each block. The first block contained a flat policy structure such that the mapping between stimuli and actions (A1, A2, etc.) was randomly assigned. The remaining four blocks all shared the same global 2^nd^-order policy structure: the shape of the stimulus indicated whether first-order rules were determined by color or texture on the current trial. In the example shown for hierarchical block 1, a circular stimulus indicated that color determined the correct action (i.e., green pairs with A1, orange pairs with A2). Hierarchical blocks included an irrelevant fourth dimension (stimulus position on screen) that is not shown here. (B) Schematic of trial design. Trials began with stimulus presentation, after which subjects had up to 2s to respond by pressing one of four buttons mapped to their right index, middle, ring, and pinky fingers. Subjects then indicated their confidence in their answer by positioning a black bar along the screen in a one-shot manner. Subjects received auditory and visual feedback following a jittered ISI.

## Results

### State-Space Model Reveals Discovery and Transfer of Global Hierarchical Structure

Trial outcomes from each block were fit with a state-space model (Figure 2A, Smith et al., 2004), and the following metrics were computed from the learning curves in each block: the (1) maximal 1^st^ derivative (mean ± within-subjects SEM: Flat, 0.006 ± 0.003; Hier 1, 0.019 ± 0.004; Hier 2, 0.024 ± 0.003; Hier 3, 0.037 ± 0.004; Hier 4, 0.037 ± 0.004, Figure 2B), and (2) maximal 2^nd^ derivative (mean ± within-subjects SEM: Flat, 0.0022 ± 0.0011; Hier 1, 0.0052 ± 0.0012; Hier 2, 0.0067 ± 0.0010; Hier 3, 0.0125 ± 0.0019; Hier 4, 0.0113 ± 0.0015, Figure 2B), and (3) the “learning trial” (mean ± within-subjects SEM: Flat, 79.29 ± 8.82; Hier 1, 48.08 ± 5.12; Hier 2, 36.83 ± 5.60; Hier 3, 33.67 ± 4.92; Hier 4, 24.12 ± 5.53, Figure 2B) (see Material and Methods for definitions).

**Figure 2.**
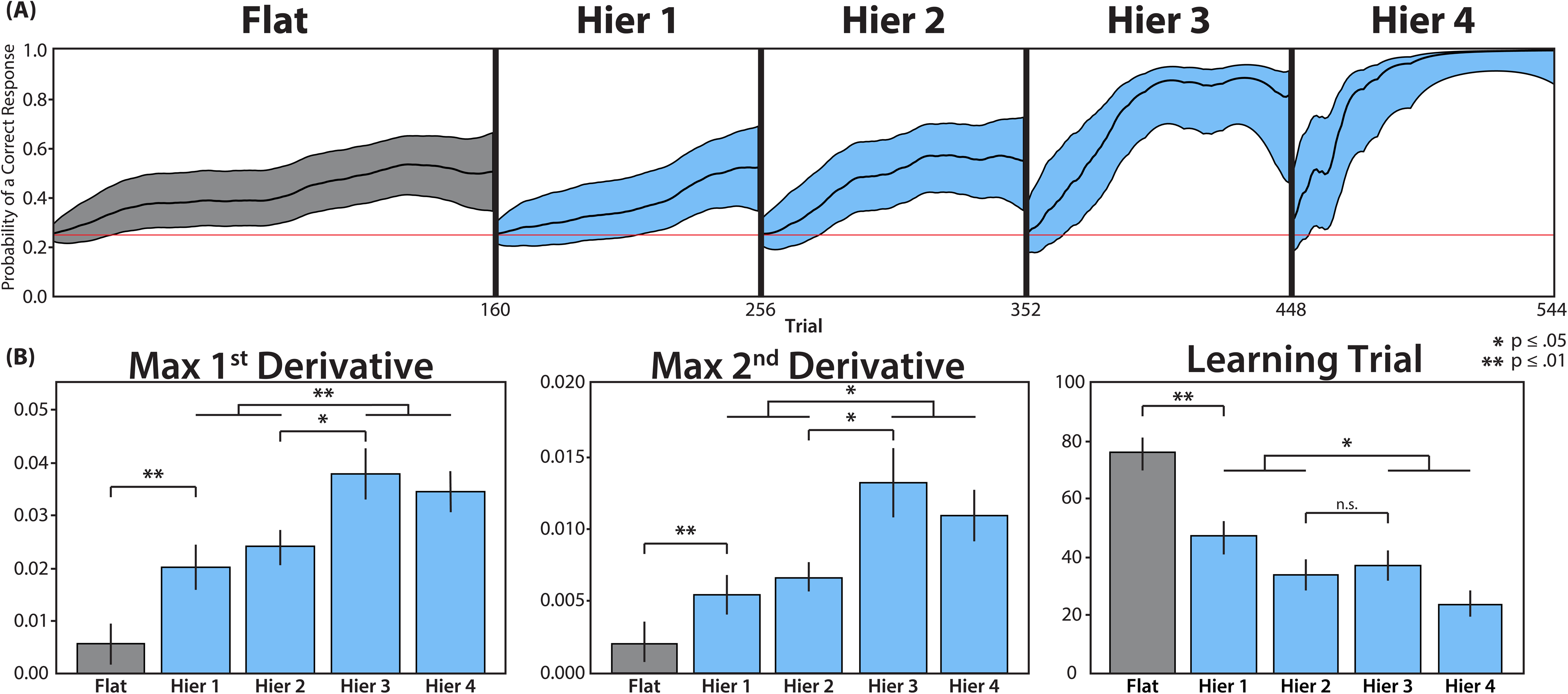
Learning Curve Results Reveal the Discovery and Transfer of the Global Hierarchical Policy Structure. (A) Output of the state-space model (Smith et al., 2004) for a representative subject. For each trial within a block, the model computes the probability of a correct response given the block’s trial outcomes. The 90% confidence interval around each trial’s estimated probability is shown in grey (Flat block) and blue (Hierarchical blocks). The red line indicates chance-level performance. B) Maximal 1^st^ and 2^nd^ derivatives and learning trial metrics derived from the model, averaged across subjects. The 1^st^ and 2^nd^ derivative metrics reveal a significant increase in learning following the 2^nd^ hierarchical block, while the mean learning trial improves more gradually across hierarchical blocks. Error bars indicate within-subjects standard error of the mean.

We first tested whether subjects acquired 2^nd^-order hierarchical rules in blocks that contained a hierarchical policy structure, which should be reflected in differences in the learning curve metrics. Compared to the flat block, learning in the first hierarchical block was more efficient (earlier learning trial: t(23) = 3.22, p = .004; Figure 2B) and showed the abrupt gains in accuracy expected from generalization of learned 2^nd^-order policy to unknown 1^st^-order rules (greater max 1^st^ derivative: t(23) = 3.30, p = 0.003; max 2^nd^ derivative: t(23) = 3.20, p = 0.004; Figure 2B). This pattern was also present when comparing learning curve metrics from the flat block to the average metrics across all hierarchical blocks: max 1^st^ derivative: t(23) = 5.74, p < 0.001; max 2^nd^ derivative: t(23) = 4.68, p < 0.001; learning trial: t(23) = 3.95, p < 0.001, Figure 2B).

We next sought to investigate the role of hierarchical structure transfer. In the first hierarchical block, subjects acquired and exploited the block-specific 2^nd^-order policy to facilitate learning relative to the flat block. Subsequently, the second hierarchical block provides the opportunity for subjects to discover the global 2^nd^-order policy structure: after acquiring the block-specific 2^nd^-order policy in the second hierarchical block, subjects can discover that the same abstract 2^nd^-order policy (i.e., shape cues color or texture) has been shared across the first two hierarchical blocks. Subjects can then transfer their learned knowledge of a global 2^nd^-order policy structure to subsequent blocks, which should greatly facilitate the acquisition of a block-specific 2^nd^-order policy (e.g. star cues color, trapezoid cues texture) and subsequently allow the subject to more rapidly resolve 1^st^-order rules within the known hierarchical structure. Thus, we predicted that successful structure transfer would result in markedly more efficient and abrupt learning following the second hierarchical block.

To test for behavioral evidence of hierarchical structure transfer, performance in hierarchical block three – where subjects can implement learned structure knowledge from the start of the block – was compared to hierarchical block two – where subjects can initially discover the global 2^nd^-order policy structure (Figure 2B). As predicted, there is a significant improvement in hierarchical learning as measured by the max 1^st^ derivative (t(23) = 2.25, p = 0.035), and max 2^nd^ derivative (t(23) = 2.23, p = 0.036). However, the learning trial metric does not show the same pattern (t(23) = 0.41, p = 0.688). This improvement is not easily explained by general practice effects: there is not a reliable change in performance metrics from the first to the second hierarchical block – when subjects can take advantage of task practice and general familiarity with the trial procedure – but must still discover the global 2^nd^-order policy structure (as assessed by all three metrics, max t = 1.28, p = 0.21). Instead, the evidence of transfer is only observed after subjects have had the opportunity to discover the global structure in the second hierarchy block.

Following discovery of the global 2^nd^-order policy structure, hierarchical knowledge transfer can facilitate learning for all subsequent blocks. Therefore, the learning metrics averaged across hierarchical blocks three and four (when the knowledge can be implemented to support learning) were compared to the average across hierarchical blocks one and two (when the knowledge has not yet been acquired). Subjects showed evidence of improved hierarchical learning in the last two hierarchical blocks versus the first two across all behavioral metrics: max 1^st^ derivative: t(23) = 2.99, p = 0.007; max 2^nd^ derivative: t(23) = 2.76, p = 0.011; learning trial: t(23) = 2.50, p = 0.020.

Lastly, we used a method previously developed to assess hierarchical learning (Badre et al., 2010) to analyze hierarchical structure learning and transfer. Instead of modeling all trials together within a block, responses to each unique stimulus were individually analyzed in order to obtain separate learning trials. Moreover, in tasks with hierarchically structured 2^nd^-ordered policy, one can conclude that a 2^nd^-order rule is completely learned if all of its subordinate 1^st^-order rules are learned above chance. Then, evidence of hierarchical structure transfer can be assessed, which should allow for faster and more complete learning of 2^nd^-order rules (mean ± within-subjects SEM: Flat, 0.17 ± 0.08; Hier 1, 0.25 ± 0.10; Hier 2, 0.50 ± 0.09; Hier 3, 1.00 ± 0.12; Hier 4, 0.92 ± 0.11). Subjects learned more 2^nd^-order rules in the hierarchical blocks than in the flat block (Z = 15.5, p < 0.001). Moreover, there was a significant increase in learned 2^nd^-order rules from the second to the third hierarchical block (Z = 4.5, p = 0.008). Lastly, subjects also learned more 2^nd^-order rules in the last two hierarchical blocks than in the first two (Z = 12.0, p < 0.001). Together, these results provide evidence that learning and subsequently transferring the global 2^nd^-order policy structure supports more efficient hierarchical learning, over and above the expected level of hierarchical learning if the hierarchical policy must be re-learned on every block.

### Mixture of Experts Model Reveals Transfer of Specific Hierarchical Structure

Although learning rate metrics derived from the state-space model allow us to characterize how learning changes across blocks, they do not provide information about *why* learning may have changed. We theorized that subjects discovered the specific 2^nd^-order policy that was globally persistent across blocks. When learning the rules for a new block, this knowledge should encourage subjects to test the hypothesis that shape determines 2^nd^-order policy. In turn, this would enhance learning by biasing their attention toward the relevance of the shape dimension, and away from the color and texture dimensions. As an alternative explanation, subjects might have discovered that the presence of hierarchical policy, in general, was persistent across blocks: one dimension cues the relevant 1^st^-order dimensions. When learning the rules for a new block, this knowledge should encourage subjects to test the hypothesis that a 2^nd^-order policy exists. This knowledge could enhance learning by biasing their attention towards the relevance of 2^nd^-order policies, in general, versus a flat policy. Because the state-space model cannot distinguish these two explanations, we used a hybrid Bayesian-reinforcement learning Mixture of Experts (MoE) model to infer the latent hypothesis states of each subject during the learning process (Frank & Badre, 2012). This approach allows us to probe the underlying cognitive mechanisms that support transfer by estimating how specific hypotheses regarding hierarchical task structure were being attended and transferred across blocks (see Materials and Methods for details).

The MoE model was employed to derive attention measures for four modeled “experts” each associated with a specific hypothesis. The first measure indexes the attention subjects place on the specific hypothesis that the shape dimension forms the top of the 2^nd^-order policy and cues subordinate 1^st^-order rules based on either color or texture (referred to as “attention to the hierarchical shape expert”). The second and third measures index the attention placed on the specific hypotheses that the color or texture dimensions, respectively, form the top of the hierarchy. The fourth measure indexes the attention subjects place on the general hypothesis that hierarchical structure, in the form of any 2^nd^-order policy, exists in the block compared to a flat policy (referred to as “attention to hierarchy”). The attention to hierarchy measure does not discern between which dimension sits atop the hierarchy, in contrast to the other three measures. In order to characterize what knowledge is being transferred from the previous block, we focus on the model estimates for these measures that capture the state of the subject prior to encountering the first trial of the block (i.e. at the “0^th^ trial”). Therefore, a discrimination can be made between whether a subject is transferring a hypothesis regarding a specific 2^nd^-order policy (attention to the hierarchical shape expert), compared to a general hypothesis regarding the presence of 2^nd^-order policy (attention to hierarchy), at the start of the block.

First, in order to determine whether subjects discover the global 2^nd^-order policy that is persistent across blocks and then test the hypothesis that this policy applies to subsequent blocks, the attention to the hierarchical shape expert was analyzed across blocks (mean ± within-subjects SEM: Flat, 0.30 ± 0.04; Hier 1, 0.58 ± 0.05; Hier 2, 0.58 ± 0.07; Hier 3, 0.77 ± 0.05; Hier 4, 0.73 ± 0.06, Figure 4A). In line with our predictions, subjects’ attention to the hierarchical shape expert at the start of the block increases from the second to the third hierarchical block (t(23) = 2.08, p = 0.049, Figure 3A), after they have had the opportunity to discover the global 2^nd^-order policy structure. Moreover, because this knowledge can inform the hypotheses for all subsequent blocks, attention to the hierarchical shape expert is greater at the start of hierarchical blocks three and four than at the start of the first two hierarchical blocks (t(23) = 2.64, p = 0.015). Although specific statistical predictions regarding attention to the hierarchical color and texture experts were not made, attention to these experts should generally be diminished when attention is biased in favor of the hierarchical shape expert. Indeed, attention to the color and texture experts is qualitatively low in the hierarchical blocks (Figure 3B, C).

**Figure 3.**
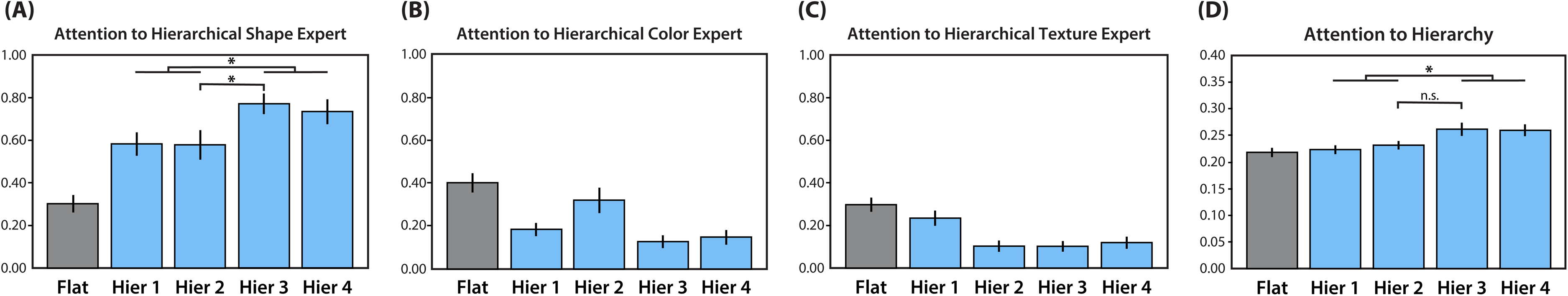
Trial Responses Reveal Transfer of Specific Hierarchical Structure. Mixture of Experts model weights for Attention to the Hierarchical (A) Shape, (B) Color, and (C) Texture Experts, as well as (D) Attention to Hierarchy, at the beginning of the Flat (gray) and four Hierarchical (blue) blocks. Each expert corresponds to a latent hypothesis regarding the hierarchical task structure that a subject might hold at the beginning of each block. Following the second hierarchical block, there is a significant increase in attention for the expert that corresponds to the global 2^nd^-order policy: shape cues color or texture (A), while attention to general hierarchical structure changes more gradually across blocks (D). Error bars represent within-subjects standard error. Significance is assessed at p < 0.05.

**Figure 4.**
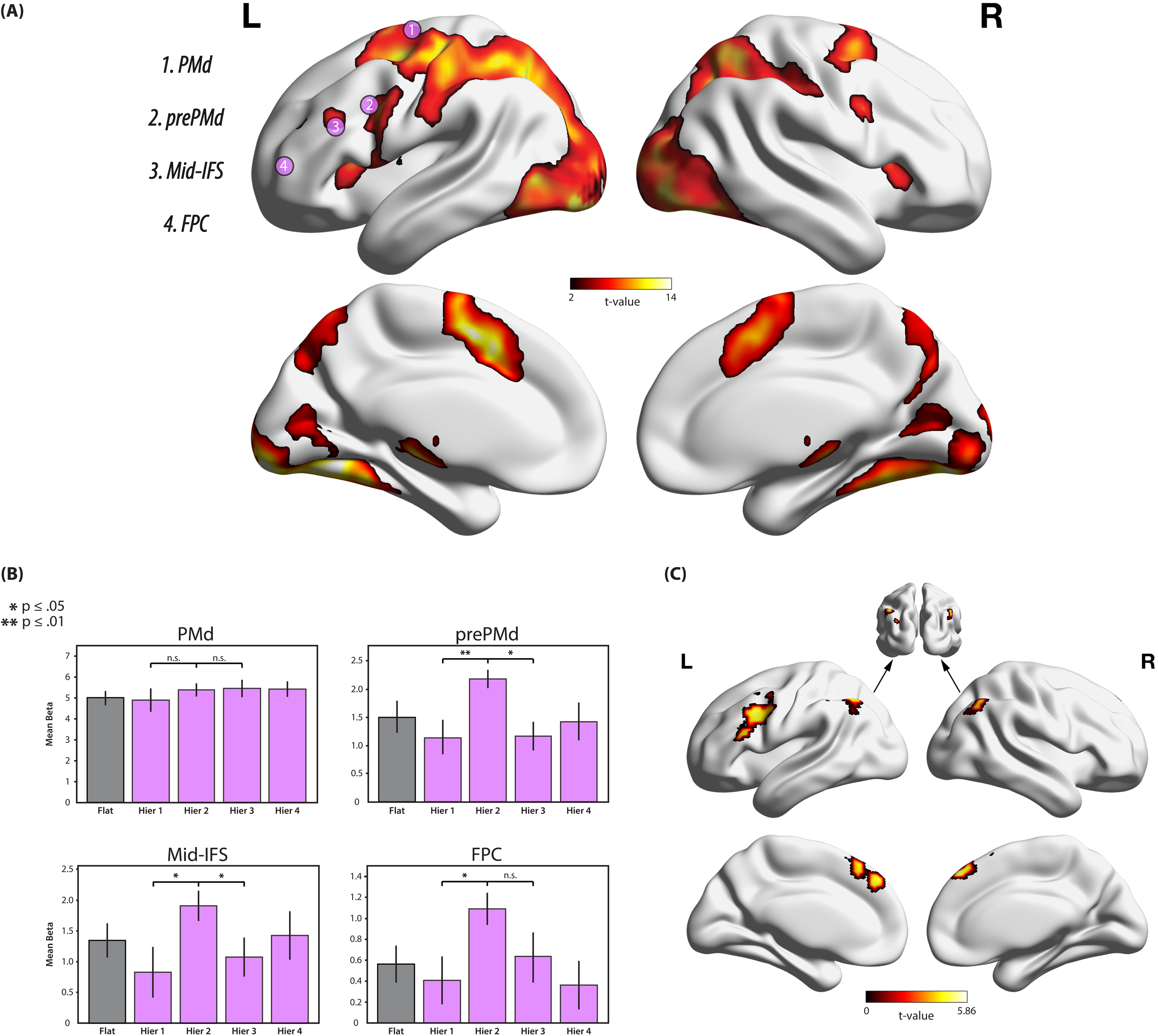
Frontal Regions Linked to Discovery of Global Hierarchical Policy Structure. (A) Group-level activity across all blocks during the stimulus response phase on correct trials only. The overlaid numbered pink circles indicate the position of each of the four lateral frontal cortex ROIs. (B) ROI analyses for the regions shown in A. The mean beta coefficients from the stimulus response phase show elevated activity during the second hierarchical block (versus the first and third hierarchical blocks) in all regions except PMd. Error bars indicate within-subject standard error. (C) Activity during the search and discovery of global hierarchical structure during the second hierarchical block shown by the contrast of Hier 2 > Hier 1 + Hier 3. All activity maps are cluster-corrected to a family-wise error rate of p < .05.

Next, we analyzed whether subjects test the hypothesis that a hierarchical policy, in general, is persistent across blocks (mean ± within-subjects SEM: Flat, 0.22 ± 0.01, Hier 1, 0.22 ± 0.01; Hier 2, 0.23 ± 0.01; Hier 3, 0.26 ± 0.01; Hier 4, 0.26 ± 0.01, Figure 4D). Subjects’ attention to hierarchy does not increase from the second to the third hierarchical block (t(23) = 1.70, p = 0.103, Figure 3D). However, there is a more gradual change in attention to hierarchy such that the measure increases from the first two hierarchical blocks to the last two (t(23) = 2.51, p = 0.019). Together, these results show that the improvement in hierarchical learning observed after the second hierarchical block can be explained by subjects discovering and then transferring their knowledge of the appropriate global 2^nd^-order policy structure that is persistent across all hierarchical blocks.

### Regions Linked to Discovery of Global Hierarchical Structure

First, a whole-brain univariate contrast of activity during the stimulus response phase on correct trials across all blocks compared to baseline was performed (p = 0.05 cluster-corrected, extent threshold: 29516 voxels, Figure 4A). The resultant map is consistent with those seen in previous hierarchical reinforcement learning studies (Badre et al., 2010). The task recruited regions along the lateral frontal cortex associated with hierarchical task performance (Badre & D’Esposito, 2007; Koechlin, 2003), as well as parietal cortex, and more specifically the intraparietal sulcus, anterior insula, mid-cingulate cortex, occipital lobe, thalamus, and medial temporal lobe.

To address which brain regions supported searching for and discovering the global 2^nd^-order policy structure, we first focused on regions in left lateral frontal cortex that support the learning and execution of hierarchical control policies: the dorsal premotor cortex (PMd), pre-dorsal premotor cortex (prePMd), mid-inferior frontal sulcus (Mid-IFS), and frontal polar cortex (FPC) (Badre et al., 2010, Figure 4A). In their original work, Badre and colleagues (2007) discovered that PMd resolved competition between 1^st^-order rules regarding motor response options, prePMd resolved competition between 2^nd^-order rules relating one stimulus feature to another (e.g. for squares, red cues action 1 while blue cues action 2), Mid-IFS resolved competition between 3^rd^-order rules, and FPC resolved competition between 4^th^-order task contexts. Moreover, activity in these regions has been associated with the search for a specific hierarchical policy within a task block (Badre et al., 2010). However, it remains unknown whether these same regions also support the learning of a more abstract, global hierarchical structure that facilities learning the specific hierarchical policies within each block.

The behavioral results demonstrate that subjects were able to both learn block-specific hierarchical policies, as well as search for and discover the global hierarchical policy structure during the second hierarchical block. To identify activity in the frontal cortex that is related to discovering the global structure, over and above activity associated with learning a block-specific hierarchical policy, activity in the second hierarchical block relative to the first hierarchical block was assessed (mean ± within-subjects SEM: PMd, Hier 1, 4.90 ± 0.56, Hier 2, 5.39 ± 0.31; prePMd, Hier 1, 1.14 ± 0.31, Hier 2, 2.17 ± 0.17, Mid-IFS, Hier 1, 0.94 ± 0.48, Hier 2, 2.16 ± 0.25; FPC, Hier 1, 0.40 ± 0.24, Hier 2, 1.09 ± 0.15, Figure 4B). With the exception of PMd (t(18) = 0.72, p = 0.483), activity across the lateral frontal cortex regions is greater in the second block compared to the first (prePMd (18) = 3.48, p = 0.003); Mid-IFS (t(18) = 2.10, p = 0.050); FPC t(18) = 2.19, p = 0.042). Next, activity in the second hierarchical block was compared to the third hierarchical block, where subjects no longer need to search for structure and can instead implement their transferred structure knowledge from the second hierarchical block (mean ± within-subjects SEM: PMd, Hier 2, 5.39 ± 0.31, Hier 3, 5.45 ± 0.41; prePMd, Hier 2, 2.17 ± 0.17, Hier 3, 1.16 ± 0.25; Mid-IFS, Hier 2, 2.16 ± 0.25, Hier 3, 1.21 ± 0.36; FPC, Hier 2, 1.09 ± 0.15, Hier 3, 0.62 ± 0.26, Figure 4B). Again, activity in prePMd (t(18) = 2.80, p = 0.012) and Mid-IFS (t(18) = 2.12, p = 0.048) is greater in the second hierarchical block. Activity is also numerically greater in FPC (t(18) = 1.52, p = 0.147), but not statistically significant. Lastly, activity in PMd did not differ across the blocks (t(18) = 0.12, p = 0.904). Because the activity in rostral regions of frontal cortex is elevated in the second hierarchical block relative to both the preceding and proceeding blocks, the observed results are likely due to a process that is preferentially engaged in the second hierarchical block, as opposed to a process that continuously evolves over time such as effects related to time on task or practice.

Whole-brain voxelwise analysis were performed to identify other regions that were recruited when subjects search for the global hierarchical policy structure by contrasting activity in the second hierarchical block to the average of the first and third hierarchical blocks (Figure 4C). This contrast revealed activity that overlapped with the left prePMd and Mid-IFS ROIs. Activity was also found in the dorsal anterior cingulate, the left inferior parietal lobule (IPL) and intraparietal sulcus (IPS), and the right IPL. The location of these lateral frontal and parietal regions overlap with a set of regions referred to as the “FP network” (Dosenbach et al., 2007).

### Regions Linked to Transfer of Global Hierarchical Structure

Next, we determined if activity in the lateral frontal ROIs predicts behavioral transfer, which was indexed by more abrupt hierarchical learning in the blocks that follow discovery of the global hierarchical structure (see Materials and Methods for definitions and details). Different lateral frontal cortex regions could support transfer of the global hierarchical policy structure. For example, prePMd could support transfer of 2^nd^-order policy by means of a more efficient resolution of competition between competing within-block 2^nd^-order rules. Alternatively, if transfer is an additional third level in the policy hierarchy (i.e. the task block contextualizes 2^nd^-order rules associated with the shape dimension), then Mid-IFS (e.g. the region associated with policy abstraction one level greater than that being transferred) could support transferring learned structure. Lastly, FPC activity could support transfer, as structure transfer may be a form of extended temporal contextualization, or episodic control, that biases task representations across multiple blocks. Knowledge of the shape dimension’s position in the hierarchy may take the role of a schema and thus recruit FPC to support the accommodation and contextualization of new information within this framework.

To test these predictions, correlations between the mean activity in each frontal ROI from blocks where behavioral transfer could occur (i.e. hierarchical blocks three and four), and the behavioral metric of transfer for each subject were performed (Figure 5A). Activity in prePMd (r = – 0.02, p = 0.937), Mid-IFS (r = 0.15, p = 0.528), and FPC (r = 0.18, p = 0.633) did not reliably correlate with behavioral transfer. However, activity in the most caudal frontal region, PMd, did reliably correlate with behavioral (r = 0.65, p = 0.002).

**Figure 5.**
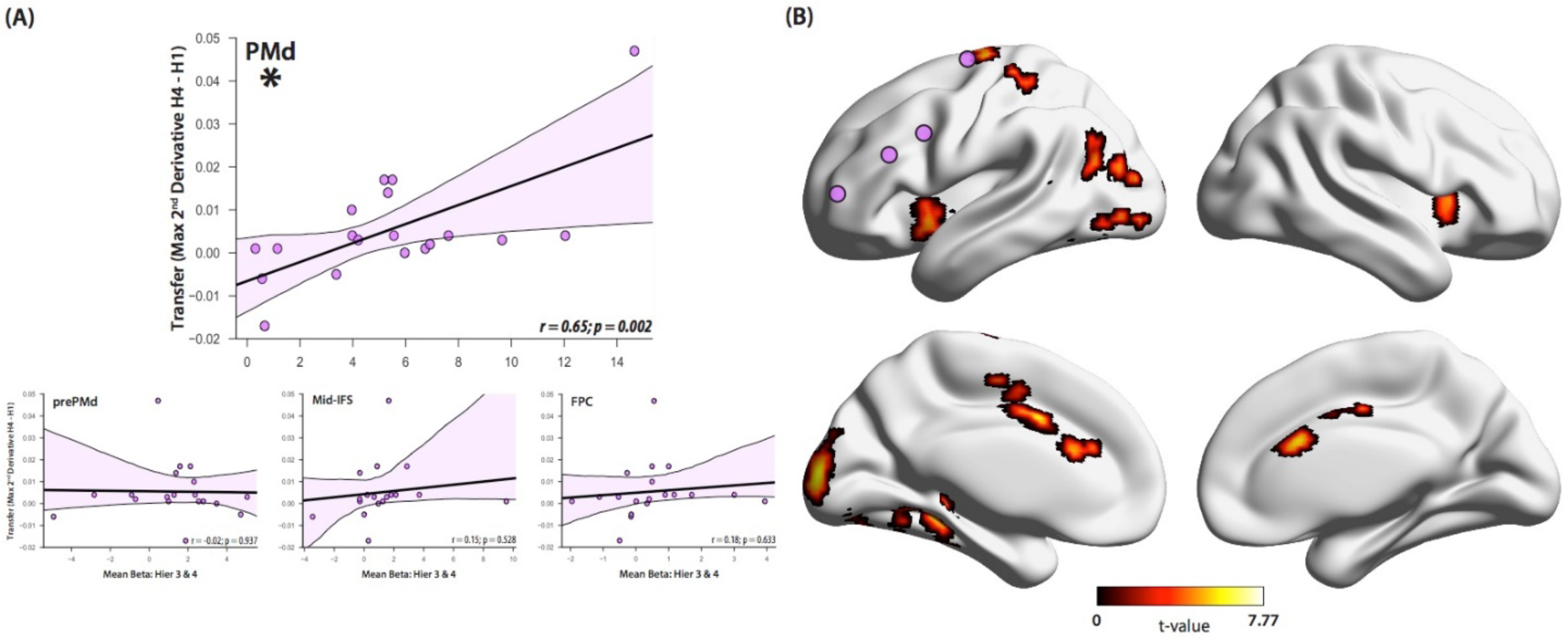
Caudal Lateral Frontal Cortex Activity Correlates with Structure Transfer. (A) Correlations between behavioral transfer and activity in the left lateral frontal cortex ROIs following discovery of the global hierarchical policy structure. Only activity in PMd was significantly correlated with individual differences in transfer. (B) Whole-brain analysis of regions for which stimulus response phase activity following discovery of the global hierarchical structure correlates with behavioral transfer (cluster-corrected at FWE p < 0.05). Overlaid pink circles indicate the position of the lateral frontal cortex ROIs.

To further identify which regions correlate with behavioral transfer in blocks following the discovery of the global hierarchical policy structure, a whole-brain analysis was performed using the degree of behavioral transfer as a parametric modulator of the mean stimulus response phase activity in the third and fourth hierarchical blocks (p = 0.05 cluster-corrected, extent threshold: 109 voxels, Figure 5B). PMd activity (overlapping with our ROI) – in accord with the previous ROI analyses – as well as bilateral anterior insula / frontal operculum, anterior cingulate cortex, left lateral occipital cortex, and left medial temporal cortex correlated with behavioral transfer.

Anterior insula and dorsal anterior cingulate cortex correspond to the “core” regions of the putative cingulo-opercular network commonly found in tasks requiring cognitive control (“CO network”, Dosenbach et al., 2008; Sadaghiani & Kleinschmidt, 2016). To further characterize the relationship between transfer and these brain regions, we tested whether this relationship was present when these two regions (and others in the “CO” network) were defined based on a previous meta-analysis of cognitive control tasks (Dosenbach et al., 2007; Figure 6A). This analysis confirmed a significant relationship between activity averaged across all cingulo-opercular regions and behavioral transfer (r = 0.57, p = 0.011, Figure 6B). Since the CO network was chosen for further analysis based on the observation of insular and cingulate activity in our whole-brain analysis, any ROI analyses that include these “core” nodes may be biased by circularity (Vul, Harris, Winkielman, & Pashler, 2009). To rule out this possibility, a separate analysis was performed that included only the canonical CO network nodes outside the core – bilateral aPFC and thalamus – which found a significant correlation between mean ROI activity and the transfer metric (r = 0.66, p = 0.002).

**Figure 6.**
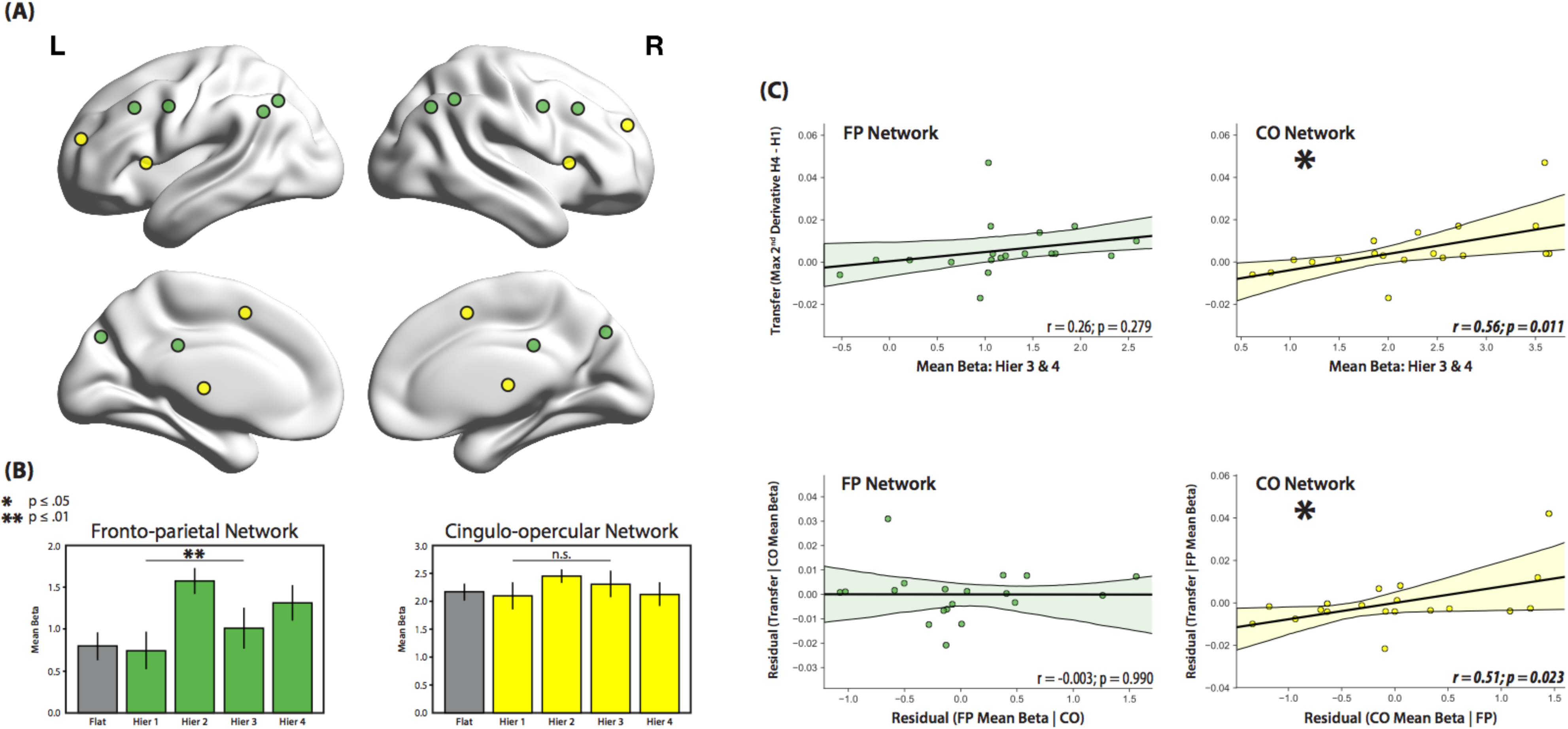
fMRI Analyses Reveal Dissociation of Behavioral Roles for FP and CO Networks. (A) Locations of regions that define the FP (green) and CO (yellow) networks defined from Dosenbach et al., 2007. FP: Bilateral frontal cortex, dorsolateral prefrontal cortex, IPL, IPS, precuneus, and midcingulate. CO: Bilateral aI/ FO, aPFC, thalamus, and dACC / msFC. (B) The contrast of the mean beta coefficients from the stimulus response phase across all respective regions in the second hierarchical block compared to the first and third blocks reveals an increase in activity during the search and discovery phase only in the FP regions. Moreover, there is a significant interaction such that the difference in activity between these blocks is greater in the FP network than in the CO network. Error bars indicate within-subject standard error. (C) Regression analyses for the FP and CO networks against behavioral transfer reveal a unique role of the CO network in structure transfer. Top row represents the correlation of transfer with each network’s mean beta coefficient from the third and fourth hierarchical blocks. Bottom row represents the partial-correlation coefficients from a multiple regression that accounts for the effects of both networks.

### Dissociation of Behavioral Roles for FP and CO Networks

The FP and CO networks have been proposed as two components of a dual-network architecture of cognitive control (Dosenbach et al., 2008), and regions in both the FP and CO networks were active during performance of our hierarchical learning task. However, these regions may support task performance by making separable behavioral contributions. To test this hypothesis, we directly compared the relationship between activity across the networks’ respective regions and (1) discovering the global hierarchical policy structure versus (2) transferring of hierarchical structure knowledge across blocks.

First, we assessed the relationship between activity in these networks and the search and discovery of hierarchical structure that occurs during the second hierarchical block. A canonical FP network was defined that included all nodes: bilateral intraparietal sulcus, bilateral frontal cortex, bilateral precuneus, bilateral inferior parietal lobule, bilateral dorsolateral prefrontal cortex, and midcingulate cortex (Figure 6A; coordinates from Dosenbach et al., 2007). Separately for the FP network and CO network ROIs, the activity for each block was estimated and a contrast was performed for the activity in the second hierarchical block versus the first and third hierarchical blocks (mean ± within-subjects SEM: FP, Hier 1, 0.74 ± 0.23, Hier 2, 1.57 ± 0.16, Hier 3, 1.01 ± 0.25; CO, Hier 1, 2.09 ± 0.25, Hier 2, 2.45 ± 0.12, Hier 3, 2.31 ± 0.24, Figure 6B). FP activity was significantly increased during the second hierarchical block (t = 3.49, p = 0.003), as expected based on the whole-brain results, whereas CO activity was not significantly different (t = 1.46, p = 0.162). Next, to formally dissociate the patterns observed across ROIs (Henson, 2006), we tested the interaction between block (second hierarchical block; average of first and third hierarchical blocks) and region (FP; CO) and found that the difference in activity between the second hierarchical block compared to the first and third blocks is significantly greater in the FP regions than in the CO regions (t = 3.37, p = 0.003).

We next assessed the relationship between activity in these networks and the transfer of hierarchical structure that occurs following discovery. Activity in the FP network in the last two hierarchical blocks was correlated with the behavioral transfer metric and no significant relationship was found (r = 0.26, p = 0.279; Figure 6C). In contrast, there was a significant correlation between activity in the CO network and the behavioral transfer metric (Figure 6C). To test whether the CO network is uniquely related to transfer, both the CO network and FP network activity were included in a multiple regression with behavioral transfer as the dependent variable, as this approach controls for any shared contribution made by both networks. This analysis revealed that only activity in the CO network reliably predicts transfer (CO network: r = 0.52, p = 0.023; FP network: r = − 0.003, p = 0.990, Figure 6B). Collectively, these findings demonstrate a clear dissociation: the regions of the FP network are more involved in the search and discovery of hierarchical structure, whereas the regions of the CO network are uniquely involved in the transfer of hierarchical structure knowledge across blocks.

## Discussion

In this study, subjects were able to efficiently discover and exploit abstract structure during a hierarchical reinforcement learning task. Specifically, during the task, subjects rapidly discovered and generalized an embedded global task structure to subsequent novel task blocks. Moreover, this generalization was supported by an increase in subjects’ awareness of the specific global hierarchical structure at the start of a new task block. The fMRI data revealed that multiple left lateral frontal regions were involved during performance of this task (prePMd, Mid-IFS, and FPC). In addition, regions within a frontal-parietal network were involved in the initial discovery of the global hierarchical policy structure. In contrast, PMd and regions within a cingulo-opercular network were involved in the transfer and implementation of this global hierarchical structure.

Our findings contribute to our understanding of the human ability to discover structure in the environment. Previous work on structure learning in the context of hierarchical reinforcement learning (Collins et al., 2014; Collins & Frank, 2013, 2016) has shown that subjects tend to build generalizable structures that allow for components of the stimulus (e.g. shape) to act as a higher-order context that cues rules based on other features of the stimulus (e.g. color). However, in contrast to previous work where stimulus-response groupings could be directly transferred, our task design prevented subjects from directly transferring action mappings across task blocks. Instead of discovering structure that immediately informed action, such as learning one of the block-specific hierarchical policies, our subjects discovered structure that informed subordinate task-set policies, as evidenced by more rapid learning in hierarchical blocks after discovering the global hierarchical structure. Moreover, when a MoE model was used to derive an estimate of subjects’ attention to the hierarchical shape rule at the start of the third hierarchical block, the model-derived estimate was greater than at the start of the second hierarchical block, indicating that subjects transferred and immediately applied their structural knowledge following discovery in the second hierarchical block. This demonstrates that subjects are capable of learning a higher-order representation between stimulus dimensions that can abstract away from the groupings of specific response pairings, and that subjects can then transfer this knowledge to new contexts to accommodate novel information.

Our brain imaging findings have implications for understanding the functional organization of the frontal cortex in support of hierarchical learning. The lateral frontal cortex is recruited for both the learning and execution of hierarchical rules (Badre & D’Esposito, 2007; Badre et al., 2010; Badre & Nee, 2018; Collins et al., 2014; Koechlin, 2003; Nee & D’Esposito, 2016), with recruitment of more rostral frontal regions during processing of higher levels of policy abstraction. In addition, patients with lateral frontal cortex lesions exhibit asymmetric behavioral impairments: caudal lesions impair both concrete and abstract cognitive control task performance, while rostral lesions only impair abstract task performance (Badre, Hoffman, Cooney, & D’Esposito, 2009). In tasks where hierarchical rules had to be implicitly learned, different lateral frontal regions are simultaneously involved in the search for hierarchical policy within a single block (Badre et al., 2010). However, patients with pre-PMd lesions are impaired at learning the full 2^nd^-order policy, but not the subordinate 1^st^-order rules (Kayser & D’Esposito, 2013). This asymmetric functional deficit is evidence of the hierarchical organization of functions associated with these lateral frontal regions. Our study extends these findings by demonstrating that frontal cortex is involved in the search for a global hierarchical policy structure, beyond that of the block-specific 2^nd^-order policies, when evidence of its presence is first available (i.e. hierarchical block 2). We conclude that the same hierarchical frontal cortex organization used to execute policy rules, as well as search for hierarchical relationships of varying complexity within the moment (i.e. block-specific policies), is also involved in the search for hierarchical relationships across putatively distinct task contexts.

In our study, only activity in the most caudal region (PMd) correlated with transfer and implementation of global hierarchical structure, which was defined as the change in the maximum 2^nd^-derivative across blocks. The maximum 2^nd^-derivative captures the initial rise of the learning curve, which indicates the transition from searching for higher-order rules to the resolution of 1^st^-order rules. In other words, subjects are transitioning from a phase of the task where the search space of possible structures is large to one where it has become well-defined and narrow. With conflict of the 2^nd^-order policy resolved, all that remains is the resolution of 1^st^-order rules, a process linked to PMd function. Subjects who resolve the 2^nd^-order conflict more rapidly can then rely primarily on processes associated with PMd (i.e., linking specific colors and textures to motor responses) for the remainder of the block, therefore facilitating performance.

Together with previous work, the current findings suggest a sophisticated coordination between motor control, rule implementation, rule discovery, and rule generalization in the service of hierarchical control, where each function incorporates knowledge of both the immediate setting (i.e. task block) and overall environment (i.e. global hierarchical policy structure across all hierarchical task blocks). In simple tasks lacking contextual elements, caudal premotor regions likely resolve response competition without influence from superordinate rostral frontal regions. However, in tasks for which contextual information must be considered (e.g., abstracted hierarchical policy), rostral premotor and mid dorsolateral regions are likely recruited to exert control over sensory-motor conflict in more caudal premotor regions (Badre et al., 2009; Kayser & D’Esposito, 2013). In settings where actions and rules are being learned, these contextual influences are likely being tested and updated via cortico-striatal interactions in response to feedback signals from the task (Badre & Frank, 2012; Frank & Badre, 2012). Thus, when a subject discovers and transfers global structure, knowledge of this structure works to restrict the search space of potential hypotheses, resulting in selective recruitment of regions along the rostrocaudal gradient to those involved in representing the generalized structure known by the individual. Thus, multiple regions across lateral frontal cortex can be involved in the process of structure transfer, but specifically only those regions along the gradient necessary for the resolution of the remaining unresolved block-specific rules.

Several cortical and subcortical regions outside the lateral frontal cortex that were associated with behavioral transfer were also identified. Specifically, subjects with greater levels of activity in regions comprising the CO network learned the block-specific hierarchical policies faster following discovery of the global hierarchical structure compared to before. Critically, this association was not found in regions comprising the FP network, suggesting that CO network activity is specifically related to the manner in which subjects maintain and implement the learned structure. Alternatively, it is possible CO activity is increased in subjects who are more engaged and attentive to the task (Sadaghiani & Kleinschmidt, 2016). However, we favor the former interpretation because our metric of transfer indexes a difference between performance in the first, compared to final, hierarchical block, and is thus insensitive to differences between subjects who perform poorly in both phases (when it could be assumed that subjects are failing to pay attention to the current task), and those who perform exceedingly well in both phases (when it is likely that attentional engagement is greatest).

Previous work has implicated the CO network in both “task-set maintenance” (Dosenbach et al., 2008, 2007, 2006), broadly defined as the configuration of control signals required to perform any type of task, and “tonic alertness” (Sadaghiani & D’Esposito, 2015; Sadaghiani & Kleinschmidt, 2016; Sadaghiani et al., 2010), or the user-driven sustained control necessary to remain prepared to process incoming information. Task-set maintenance requires that a specific structure be known to the individual – that which defines successful performance of the task – whereas tonic alertness precludes any need for a specific structural representation of the task as alertness takes the role of “nonselective disengagement” (Sadaghiani & Kleinschmidt, 2016). Thus, our findings are more consistent with a role of the CO network in task-set maintenance, although a role in tonic alertness during our task cannot be ruled out.

Whereas the CO network was uniquely related to transfer, the FP network was selectively involved in the search and discovery of the global hierarchical policy structure. Our findings suggest that the FP network is not only involved in the representation and integration of current task rules and response mappings, but also in the integration of previous task-relevant components. The integration of this information would likely allow for complex structured relationships to be discovered across blocks. Recent studies have discovered that tasks requiring varying levels of cognitive control recruit regions along a caudal-rostral gradient in parietal cortex in a similar fashion to that found in lateral frontal cortex (Choi, Drayna, & Badre, 2018). Moreover, regions along both gradients showed mirroring patterns of functional connectivity with striatal sites, in line with models of hierarchical rule learning, and task set creation and implementation (Badre & Frank, 2012; Collins & Frank, 2013). In line with this recent evidence, the present results implicate a system of parallel and distributed hierarchical gradients across frontal and parietal cortex that supports the search and discovery of structure of varying complexity within and across task blocks.

## Materials and Methods

### Human Subject Details

Thirty-two healthy right-handed subjects (range: 18 – 29 years; mean = 19.63; SD = 2.54; 20 females) with normal or corrected-to-normal vision participated in the study at the University of California – Berkeley. Target sample size was based on prior relevant literature (Badre et al., 2010; Collins & Frank, 2016; Nee & D’Esposito 2016). Eight subjects were excluded from all behavioral analyses (four subjects failed to complete the entire session, two subjects did not follow the instructions, and two subjects exhibited sub-threshold behavioral performance (no above-chance performance in any hierarchical block)). Five additional subjects were excluded from all fMRI analyses (one subject due to above-threshold in-scanner motion (>2.5mm in X, Y, or Z across all blocks), one subject for atypical anatomical data and three subjects due to scanner image reconstruction failures).

All behavioral analyses presented here include data from the 24 subjects for whom we obtained a complete behavioral dataset (range: 18 – 24 years; mean = 19.25; SD = 1.75; 16 females). All fMRI analyses presented here include data from the 19 subjects for whom we obtained a complete behavioral and fMRI dataset (range: 18 – 24 years; mean = 19.26; SD = 1.88; 13 females). Behavioral analyses restricted to these 19 subjects are separately reported in the supplement (Figure S2). All research protocols were approved by the Committee for Protection of Human Subjects at the University of California, Berkeley. Informed and written consent was obtained from all subjects prior to participation.

### Experimental and Task Design

To investigate the discovery and transfer of abstract hierarchical structure, we designed a reinforcement learning task (inspired by Badre et al., 2010) that required learning multiple distinct rule sets that shared a common abstract hierarchical structure (hierarchical blocks) or a rule set in which there was no higher-order structure (flat block). Subjects completed one flat block and four hierarchical blocks while inside the scanner (Fig. 1). Subjects viewed stimuli that varied along three or four dimensions: shape, color, black-and-white image pattern (referred to as “texture”), and stimulus position on screen (hierarchical blocks only) (Fig. 1A). For each block, stimulus dimensions could vary between two features (e.g. Color: red/blue, Shape: square/circle, etc.), resulting in 8 unique stimuli in the flat block and 16 unique stimuli in each hierarchical block. All blocks contained unique features, and thus subjects had to learn entirely new stimulus-response mappings for each block. We assigned stimulus features to blocks by random assignment.

### Stimuli

Stimuli were generated using PsychoPy (Peirce, 2007, 2008). Colors included red, green, blue, yellow, magenta, cyan, white, maroon, black, and orange. Shapes included a circle, square, rectangle, triangle, pentagon, rhombus, trapezoid, six-sided star, oval, and tear drop. Texture images were sourced from the Normalized Brodatz Texture Database (Abdelmounaime & Dong-Chen, 2013). These images included close-up photographs of various real-world textures, such as tree rings, sand dunes, snakeskin, bubbles, etc. Subjects did not report difficulty in discriminating between textures (Fig 1A). The stimuli generally subtended ∼7.5° of visual angle. Stimulus position in the hierarchical blocks was computed along an invisible circle positioned at the center of the screen with a radius subtending ∼7.5° of visual angle. The eight locations along this circle began at 27.5° clockwise from the vertical meridian and were equally spaced by 45° increments.

### Flat Block

The flat block consisted of 20 repetitions of each stimulus for a total of 160 trials. Stimulus order was randomized within each set of 8 trials so as to restrict the range of the number of trials between stimulus repetitions. On average, each stimulus was viewed once every 8 trials, ranging from 0 to 15. Prior to the start of the block, subjects had the opportunity to view all 8 stimuli created for the upcoming block.

Trials began with the presentation of the stimulus slightly offset left of the center of the screen for a maximum of 2,000ms (Fig. 1B). Stimulus composition included a black-and-white image cropped into a specific shape with a colored border. Subjects were instructed to respond to the presentation of the stimulus by pressing one of four buttons mapped to their right index, middle, ring, and pinky fingers. Responding within 2,000ms advanced the trial to the confidence response phase. This phase began with the appearance of a vertical rectangle offset right of center with a horizontal black bar appearing either on the bottom or top of the rectangle. To indicate their confidence that their most recent response was correct (i.e. receive positive feedback at the end of the trial), subjects had 1,500ms to re-press and hold down the button they had just pressed. By re-pressing the button, the black bar began to move away from its starting position at a constant rate until it reached the other side of the rectangle, a process that lasted up to 1,250ms. Regardless of the bar’s starting position, the top of the rectangle indicated 100% confidence in their answer being correct, while the bottom of the rectangle indicated 0%. Subjects were instructed to be as precise as possible with their confidence rating. Following the release of the held-down button, or after 1,250ms, both the stimulus and confidence probe disappeared from screen. Following a variable amount of time (200ms, 1,200ms, or 2,200ms), subjects received audiovisual feedback. Correct feedback involved the presentation of the word “Correct” and “$$$$$” stacked vertically in the center of the screen, as well as a pleasant tone. Incorrect feedback contained the word “Incorrect” and an unpleasant tone. The feedback stimulus persisted for 333ms. Feedback was 100% valid. The next trial then began after a variable inter-trial interval (ITI) with a mean of 1,500ms. Given the variability of event boundaries, the amount of time not spent responding to the stimulus or reporting confidence (e.g. the subject responded to the stimulus in 850ms [1250ms left over], to the confidence probe in 1,000ms [500ms left over], and released the button after 750ms [500ms left over]: thus 2,250ms remained unused) was added to the ITI. The average trial duration was 7,783ms. The order of ITIs within a block was optimized to permit estimation of the event-related response (Dale, 1999).

We assigned two stimuli to each response option so that each button had a 25% chance of being correct on any given trial. Stimulus-response mappings were independent from one another, such that no higher-order structure was present, thus requiring each response to be learned individually. Following the final trial, mean block accuracy was presented on screen.

### Hierarchical Block

We designed the hierarchical blocks identically to the flat block with the following exceptions. (1) Stimuli now included a fourth dimension: position on screen. In each of the four hierarchical blocks, the stimulus could appear in one of two locations on screen. These locations were semi-randomly selected from 8 possible equidistant positions along an invisible aperture around the center of the screen. We assigned the positions in each block in pairs, such that each pair was offset in both the x- and y-axis so as to create as large a separation and difference as possible. Position was not included in the flat block as pilot testing indicated subjects were unable to learn above chance 16 independent stimuli across four button responses in an appropriate amount of time. (2) The number of stimulus repetitions decreased from 20 to 6, resulting in a decrease in the number of total trials from 160 to 96 per block. (3) Given the new position dimension, the confidence probe was moved to the center of the screen so as not to interfere with the stimulus. (4) The position-on-screen dimension was not included in the pre-block stimulus presentation screen in which all 8 stimuli were shown.

Lastly, and most critically, all hierarchical blocks contained a 2^nd^-order policy relationship that subjects could discover and transfer across blocks so as to facilitate their learning, instead of learning 16 independent stimulus-response mappings. Specifically, the shape dimension cued 1^st^-order rules dependent on either the colors or textures, and as a result, screen position was irrelevant. By learning and exploiting this structure the number of rules to be learned decreased to four (i.e. two rules for color, two rules for texture). The same 2^nd^-order policy relationship was maintained across blocks, in that the shape dimension always cued rules based on either color or texture dimensions.

### Instructions and Training Protocol

Prior to performing the task inside the MRI scanner, all subjects completed a training session on a desktop computer to make sure they understood the task and could perform it adequately. After obtaining experimental consent, and confirming both study and MRI scanner eligibility, subjects reviewed the instructions of the task. Along with visual aids on the computer, the experimenter described the task such that subjects knew they had to learn stimulus-response mappings across multiple task blocks, however no information was provided that could cue subjects to the hierarchical structure of the task. Subjects then practiced the confidence-reporting component of the task in a guided environment using stimuli not present in the real experiment. Subjects received guided instructions indicating which button to press and how confident they should report feeling for each practice trial. Instructed confidence levels included 0%, 15%, 35%, 50%, 65%, 85%, and 100%. Subjects needed to place the confidence bar at the appropriate location along the vertical rectangle to match the instructed confidence level across 21 practice trials (3 repetitions of each level). A 93% accuracy criterion was required to progress. Subjects had to repeat the 21-trial practice block until they met criterion. The timing of all events matched that of the real experiment.

Following completion of the confidence reporting practice, subjects then performed 24 practice trials of a flat block, using the same stimuli as before. Just as in the real task, subjects had to learn eight independent stimulus response mappings across four buttons using the feedback provided at the end of each trial. No performance criterion was included, as the goal of this practice session was to familiarize subjects with the components of the task in real time.

Upon completion of the practice session, subjects were then escorted to the MRI scanner suite and placed inside the scanner. During the acquisition of an anatomical scan (details below), subjects went through the practice instructions and confidence reporting session again so as to become accustomed to both the MRI-compatible four-button response box, and to being inside the active scanner. Subjects received compensation at a rate of $20 per hour and could earn a bonus of up to $10 based on their overall trial accuracy.

### fMRI Data Acquisition

Whole-brain imaging was performed at the Henry H. Wheeler Jr. Brain Imaging Center at UC Berkeley using a Seimens 3T Trio MRI scanner using a 32-channel head coil. Functional imaging data was acquired with a gradient-echo echo-planar pulse sequence (TR = 1,000ms, TE = 33ms, flip angle = 40°, array = 84 x 84, 52 slices, voxel size = 2.5mm isotropic). T1-weighted MP-RAGE anatomical images were collected as well (TR = 2,300ms, TE = 2.98ms, flip angle = 9°, array = 256 x 256, 160 slices, voxel size = 1mm isotropic). Subject’s head movement was restricted using foam padding. Auditory feedback was presented through in-ear headphones connected to the stimulus presentation computer. The flat block consisted of a single run of 1290 TRs, while each hierarchical block consisted of 760 TRs.

### fMRI Data Preprocessing

Preprocessing was performed using FMRIPREP (Esteban et al., 2018), a Nipype (Gorgolewski et al., 2011) based tool. Each T1w (T1-weighted) volume was corrected for INU (intensity non-uniformity) using N4BiasFieldCorrection v2.1.0 (Tustison et al., 2010) and skull-stripped using ANTs BrainExtraction. Spatial normalization to the ICBM 152 Nonlinear Asymmetrical template version 2009c was performed through nonlinear registration with the antsRegistration tool of ANTs v2.1.0 (Avants, Epstein, Grossman, & Gee, 2008), using brain-extracted versions of both T1w volume and template. Brain tissue segmentation of cerebrospinal fluid (CSF), white-matter (WM) and gray-matter (GM) was performed on the brain-extracted T1w using FSL’s fast (Zhang, Brady, & Smith, 2001). Functional data were motion corrected using FSL’s mcflirt (Jenkinson, Bannister, Brady, & Smith, 2002). This was followed by co-registration to the corresponding T1w using boundary-based registration (Greve & Fischl, 2009) with 9 degrees of freedom, using flirt (FSL). Motion correcting transformations, BOLD-to-T1w transformation, and T1w-to-template (MNI) warp were concatenated and applied in a single step using ANTs ApplyTransforms using Lanczos interpolation. Slice timing correction was not performed. Preprocessed data were spatially smoothed with an 8mm FWHM isotropic Gaussian kernel. Motion estimates used for subject exclusion were calculated using SPM’s realign function.

### Behavioral Data Analysis

Analyses of behavioral data included the use of paired t-tests with one exception. When analyzing the number of learned 2^nd^-order rules across blocks, we used Wilcoxon sign-ranked tests due to the non-parametric nature of the data (i.e. subjects could learn either zero, one, or two 2^nd^-order rules per block) and the within-subjects design of the study. In addition, the stimulus dimension of position-on-screen was fully ignored in all analyses of the data.

### State-Space Model

Trial responses were modeled with a state-space modeling approach (Smith et al., 2004) to produce learning curves. The model outputs trial-by-trial estimates of the probability of a correct response on each trial, as well as a 90% confidence interval around each estimate. Similar to (Badre et al., 2010), our analyses focused on the following metrics derived from the learning curve: (1) the trial for which the 90% confidence interval no longer included chance performance, referred to as the “learning trial”; (2) the maximal 1^st^ derivative of the learning curve, which indexes the rate of learning; (3) the maximal 2^nd^ derivative, which indexes the rate of change in one’s learning rate.

### Mixture of Experts Model

We make use of a hybrid Bayesian-reinforcement learning mixture of experts (MoE) model previously used by Frank and Badre (2012) in order to estimate subjects’ attention to various hypothesis states that we assume are being tested while subjects perform the task. Given the observed stimuli and responses, the MoE model estimates individual subjects’ attention to likely hypotheses about the relationship between context (i.e. the features of the stimulus) and action (i.e. the available button responses) in each task block. Each expert in the model represents a prediction about how a stimulus feature, or combination of features, relates to the likelihood of obtaining a reward given the motor actions available to the subject. For example, the “shape expert” could learn the likelihood of obtaining a reward based only on the shape of the stimulus. For each trial, the expert makes its prediction about what action is likely to be correct given its assigned feature, and experts who contribute accurate predictions are rewarded while experts providing unreliable predictions are not. For hierarchical experts, the model makes predictions about subordinate stimulus dimensions (i.e. color or texture) contingent on the identity of a third, superordinate dimension (i.e. color), such that weights assigned to predictions about each subordinate dimension are dynamically gated based on the feature of the superordinate dimension (i.e. blue vs. red). The MoE model also assigns attentional weights to experts that learn the overall reliability of hierarchical vs. flat predictions based on the reliability of all the hierarchical and flat experts, respectively.

For the current study, we adapted the model in order to allow for individual fits to each hierarchical expert. As the original version used a single hyperparameter across all three hierarchical experts, thus preventing the ability to estimate different initial weights, we instead modeled each hierarchical expert with a separate parameter. We also removed the decay parameter originally used to model the degree to which the current block’s attentional weights carried over into the next block. Instead, we modeled a separate set of parameters for the various experts in each block. By removing the decay parameter, and modeling each block independently, we ensure that the model is incapable of being biased by the previous block. As a result, any differences between blocks in the parameter values, as well as the computed attentional weights at the beginning of the block, are the result of that block’s data alone. In order to obtain the best fit for the data, we first modeled all subjects together (pseudo-R^2^ = 0.25 and 0.12 for the mean hierarchical block and flat block, respectively) and then used the fitted parameter values as our initialization point for the model when fitting each subject individually (mean pseudo-R^2^ = 0.33 and 0.15 for the mean hierarchical block and flat block, respectively). These pseudo-R^2^ values are similar to those reported in Frank and Badre (2012). Validation of the revised MoE model involved simulating datasets across each of the five task blocks. We used the parameter values obtained from fitting the model to the real subject data to generate simulated responses to the task. In order to draw comparisons to the human data, the simulated data was then fit to the State Space model so as to produce learning curves, which allowed for calculation of learning metrics (i.e., maximum 2^nd^ derivative). Overall, the revised MoE model was successfully able to recreate the qualitative patterns of behavior and attentional weight recovery across blocks seen in the human data (Figure S1).

### Mixture of Experts model validation procedure

Our initial validation step was to generate simulated datasets using the MoE model in order to recreate the behavioral results seen from the state-space model. We initially fit the MoE model to each subject’s set of responses in order to obtain parameter values for each block. These values were then used to generate simulated datasets. For each block, we randomly chose a subjects’ block-specific set of fitted parameter values and used these values to generate a series of responses to the same exact stimuli that were presented to the subject originally. We performed this procedure 100 times per subject per block. Learning curves were generated for each individual iteration using the state-space model, and then averaged together so as to produce a single mean simulated learning curve per subject per block. We then calculated the maximum 1^st^ and 2^nd^ derivatives for each of these mean learning curves (Fig. S1 A, B).

In order to assess the model’s ability to recover model-derived attentional weight values, we followed the procedure described above and generated 500 new datasets for each block, randomly sampling the subjects’ sets of block-specific fitted parameter values, resulting in 2,500 datasets. We then fit the model to the data in order to obtain fitted parameter values. We followed the fitting procedure as described in the main text. Specifically, we used the fitted parameter values that resulted from previously fitting the MoE model to the entire group of subjects together as the initialization point for the optimization algorithm. Using these recovered parameter values, we then computed the attentional weight values for each hierarchical expert, as well as the overall attention to hierarchy weight, on trial 0 (Fig. S1 C-H). Overall, we found a robust association between the weights computed from the real data to those from the simulated data for both attention to the hierarchical shape expert (r = 0.251, Fig. S1 C) and attention to hierarchy (r = 0.294, Fig. S1 D) on trial 0. Moreover, the pattern of mean attentional weight values qualitatively matches that seen when fitting the model to the human subject data (Fig. S1 E-H).

### Univariate fMRI Analysis

Statistical models were constructed for each subject under the assumptions of the general linear model (GLM) using SPM 12 (Statistical Parametric Mapping; www.fil.ion.ucl.ac.uk/spm). Each trial was modeled by one of two sets of five boxcar regressors: (1) a regressor for the stimulus response phase (beginning with stimulus onset and ending when a response was made), (2) a regressor with the same onset and duration as the stimulus response phase, but whose value was parametrically modulated by the subject’s reaction time to the stimulus, (3) a regressor for the confidence response phase (beginning and ending with the onset and offset, respectively, of the confidence probe), (4) a regressor with the same onset and duration as the confidence response phase, but whose value was parametrically modulated by the reported confidence level, (5) a regressor for the feedback phase (beginning and ending with the onset and offset, respectively, of the audiovisual feedback). In order to match the analysis approach of Badre et al., 2010, one set of regressors exclusively modeled correct trials, while the other set exclusively modeled incorrect trials. To ensure the parametrically modulated regressors only explained the variance unique to processes associated with the modulatory values (i.e. stimulus reaction time and confidence level), we orthogonalized both the modulated stimulus response phase and modulated confidence response phase regressors with respect to their respective unmodulated regressors. Next, we included three additional regressors to remove variance associated with events related to the subject failing to make a required response. Two regressors modeled stimulus and confidence response phases where no stimulus or confidence response, respectively, was made. The third regressor modeled feedback phases where “No Response” was presented. Although trials where subjects failed to indicate their level of confidence could be separated by whether the subject’s stimulus response was correct or incorrect, we chose to model these events together because we considered both events to be of no interest and thus nuisance signals. Lastly, five block regressors were included to account for run-to-run variance. In total, each block contained a theoretical maximum of fourteen regressors: some subjects had blocks where all required responses were made, and thus no regressors could be made that modeled events related to a failure to respond. Low frequency signals were removed with a 1/128 Hz high-pass filter. This first level regression thus yielded standardized regression coefficients (“betas”) for each voxel in the brain for each regressor included in the model. Linear contrasts were used to obtain subject-specific effects, which were then entered into a second-level analysis treating subjects as a random effect and comparing voxel effects against a value of zero.

Region-of-Interest (ROI) analyses supplemented the whole-brain search. ROIs were constructed with the Marsbars (Brett, Anton, Valabregue, & Poline, 2002) and wfupickatlas (Maldjian, Laurienti, Kraft, & Burdette, 2003) toolboxes in SPM12. Coordinates and sphere size for frontal cortex nodes (i.e. PMd, pre-dorsal premotor cortex (pre-PMd), mid inferior frontal sulcus (Mid-IFS), and FPC) were taken from Badre et al., (2010). Cingulo-opercular and fronto-parietal coordinates and size (i.e. CO: bilateral anterior prefrontal cortex, bilateral anterior insula / frontal operculum, bilateral thalamus, and dorsal anterior cingulate cortex / mid-superior frontal cortex; FP: bilateral intraparietal sulcus, bilateral frontal cortex (roughly BA 6), bilateral precuneus, bilateral inferior parietal lobule, bilateral dorsolateral prefrontal cortex (roughly BA 9/46), and midcingulate cortex) were taken from Dosenbach et al., (2007).

Whole-brain analyses were performed in SPM and cluster correction was performed at the family wise error rate of p = 0.05, using p = 0.001 as the cluster defining threshold. Correlations between fMRI data and behavioral data were performed using standard parametric linear regression.

### Behavioral Metrics of Transfer

To test for brain-behavior correlations that relate individual differences in transfer performance to fMRI activity, we calculated the behavioral metric of transfer based on the state-space model we employed. We computed a difference score between the fourth and the first hierarchical block so as to assess the maximum impact that hierarchical structure transfer could have on behavioral performance. Specifically, our metric of transfer came from computing the change in the state-space model’s maximum 2^nd^ derivative measure. We chose to focus on the maximum 2^nd^ derivative as it should best capture the degree to which learning accelerates once the subject determines the appropriate 1^st^-order rules associated with the known 2^nd^-order policy. Defining transfer in this manner allowed us to contrast subjects’ performance when learning a hierarchically structured task with no ability to transfer knowledge of 2^nd^-order policy to when subjects have the greatest likelihood of transferring learned 2^nd^-order policy.

For the whole-brain analysis, we defined a contrast for each subject that contrasted mean stimulus response phase activity for the third and fourth hierarchical block against baseline. At the second level, the transfer metric was used as a covariate and regressed against this contrast to identify univariate activity across individuals that was associated with differences in transfer.

## Funding Source

This work was funded by a grant from the NIH, MH63901.

## Acknowledgements

We thank Michael Frank and Anne Collins for sharing the original, and discussing appropriate revisions to, the Mixture of Experts code.

## Author Contributions

Conceptualization: A.E. and J.S.; Methodology: A.E. and J.S.; Software: A.E. and J.S.; Formal Analysis: A.E. and J.S.; Investigation: A.E.; Resources: M.D.; Writing – Original Draft: A.E.; Writing – Review & Editing: A.E., J.S., and M.D.; Visualization: A.E.; Supervision: J.S. and M.D.; Project Administration: A.E.; Funding Acquisition: M.D.

## Competing Interests

The authors declare no competing interests.

## Supplemental Information

**Figure S1.**
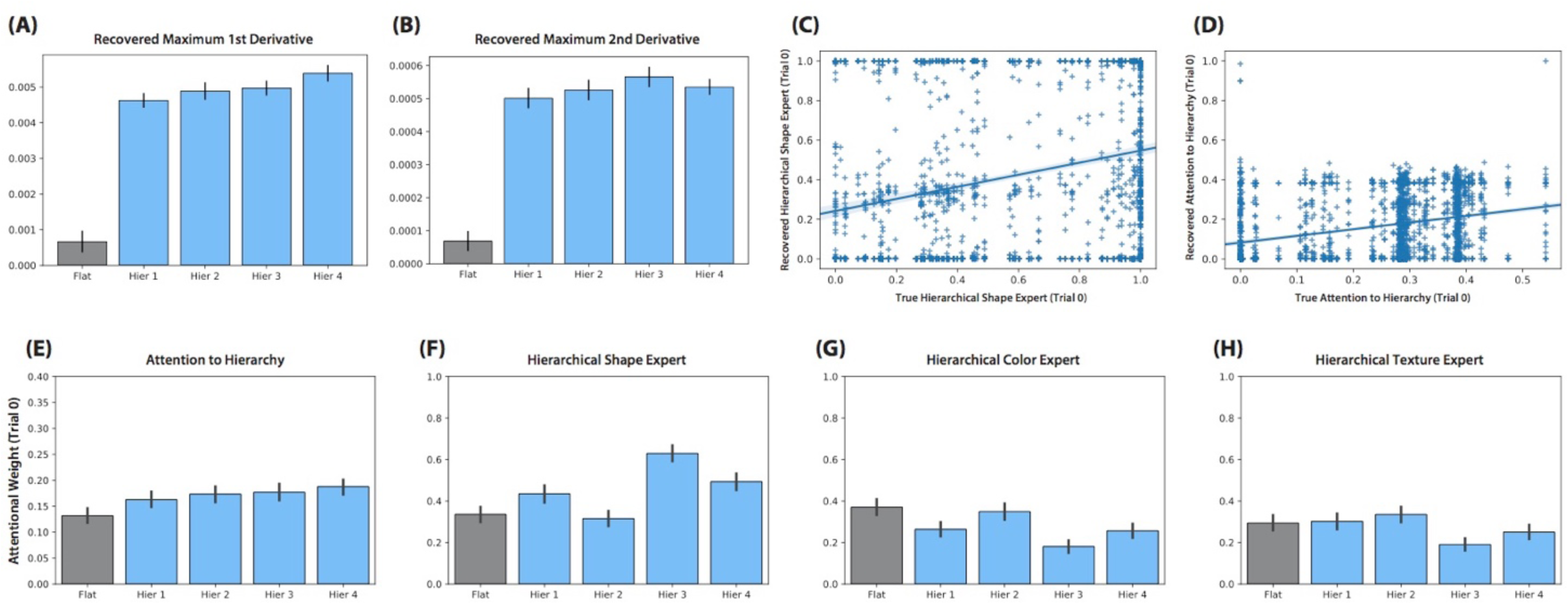
Model validation results for the Mixture of Experts model. (A) The mean maximal 1^st^ and (B) 2^nd^ derivatives from the mean state-space model learning curves. (C) The relationship between the attentional weight to the hierarchical shape expert on trial 0 computed from the real subject data and the value computed from the simulated data. Given that there existed a positive relationship between the real and simulated results, shown here are the 2,500 data points collapsed across all five blocks, along with a least square best-fit line to represent the linear relationship. (D) Same as (C) but for the attention to hierarchy weights. (E) Mean attention to hierarchy on trial 0 computed from the simulated data is shown for each task block. (F-G) Same as (E), but for the (F) hierarchical shape, (G) hierarchical color, and (H) hierarchical texture experts. Y-axis values for (E-H) match those seen in Figure 2 in the main text. All error bars are within-subject standard error of the mean.

**Figure S2.**
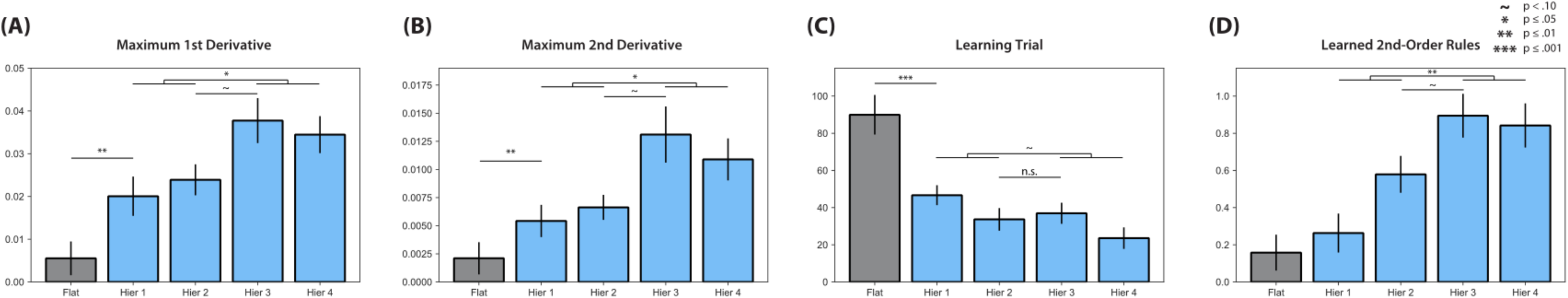
Analysis of state-space learning curve data in cohort of 19 subjects eligible for fMRI analyses. (A) The mean maximal 1^st^ and (B) 2^nd^ derivative of the learning curve across the five task blocks. In both cases, learning was faster in the first hierarchical block than in the flat block (max 1^st^ derivative: t(18) = 3.08, p = 0.007; max 2^nd^ derivative: t(18) = 2.97, p = 0.008). The increase in learning rate from the second to the third hierarchical block almost reached significance for both measures (max 1^st^ derivative: t(18) = 1.86, p = 0.080; max 2^nd^ derivative: t(18) = 1.98, p = 0.063). Overall, learning in the last two hierarchical blocks occurred more rapidly than in the first two (max 1^st^ derivative: t(18) = 2.29, p = 0.034; max 2^nd^ derivative: t(18) = 2.25, p = 0.037). (C) Mean learning trial across the task blocks. The learning trial occurred earlier in the first hierarchical block than in the flat block (t(18) = 4.28, p < 0.001). The learning trial tended to occur earlier in the final two hierarchical blocks than in the first two (t(18) = 1.76, p = 0.096), but was no different between the second and third hierarchical blocks (t(18) = 0.40, p = 0.696). (D) The mean number of learned 2^nd^-order rules across the task blocks. Subjects tended to learn more 2^nd^-order rules in the third hierarchical block than in the second hierarchical block (Z = 3.50, p = 0.058). Overall, subjects learned more 2^nd^-order rules in the final two hierarchical blocks than in the first two (Z = 5.50, p = 0.002).

